# Lateral hypothalamic GABAergic projections to the dorsal pons and lateral preoptic area in feeding, predation, and reinforcement

**DOI:** 10.64898/2026.08.13.744663

**Authors:** Yi Huang, Will Fan, Olivia Knuth, Alexander C. Jackson, Natale R. Sciolino

## Abstract

Lateral hypothalamic GABAergic (LHA^GABA^) neurons regulate arousal, feeding, and reward-related behaviors, but how their downstream projections coordinate motivated behaviors across domains remains incompletely defined. Our histological analyses revealed that LHA^GABA^ fibers were distributed across the dorsal pons (DP) subregions, including the peri-locus coeruleus, laterodorsal tegmentum, and Barrington’s nucleus, and extend throughout the lateral preoptic area (LPO), thereby refining existing anatomical descriptions. We then used optogenetics to systematically compare the effects of activating LHA^GABA^ somata and their projections to the DP and LPO across assays of feeding, non-food-directed gnawing, predatory behavior, real-time place preference, and operant self-stimulation. In sated mice, optogenetic activation of LHA^GABA^ somata or their terminals in the DP or LPO increased caloric food intake, whereas non-caloric cellulose intake was minimally affected during terminal stimulation. Across conditions, activation increased gnawing and shredding of non-food objects while reducing inactivity. In cricket hunting, stimulation increased cricket killing and consumption relative to controls. Similarly, all stimulation conditions supported positive-valence and reinforcement-related responding, as indicated by real-time place preference and operant self-stimulation. Together, these results provide new functional evidence that activation of LHA^GABA^ somata and projections to both the DP and LPO recruit largely overlapping behavioral responses across feeding, non-food behavior, predatory hunting, and reinforcement-related assays. These findings support a distributed hypothalamic output architecture in which major ascending and descending LHA^GABA^ pathways contribute to a shared motivational repertoire rather than wholly discrete behavioral functions.

## INTRODUCTION

The lateral hypothalamic area (LHA) is a key regulator of arousal, appetitive and consummatory behavior. Positioned at the intersection of neural and humoral signaling systems, the LHA coordinates programs that maintain physiological homeostasis and survival-related motivated behavior^1–9^. Despite this central role, the circuit mechanisms through which the LHA organizes motivated behavior remain incompletely understood. A major population implicated in these functions is its GABAergic neuronal population.

Lateral hypothalamic GABAergic neurons (LHA^GABA^), identified by expression of key components required for GABA synthesis and vesicular release, including GAD65, GAD67, and VGAT (SLC32A1, encoded by *Slc32a1*), comprise a substantial population of LHA neurons^4,8,10–13^. Single-cell transcriptomic and spatial mapping studies have further revealed marked molecular heterogeneity and spatio-molecular organization within the LHA, including among its GABAergic neurons^14–16^. Consistent with this diversity, LHA^GABA^ neurons regulate a broad range of behaviors. Optogenetic or chemogenetic stimulation of LHA^GABA^ neurons enhances feeding and reward-related behaviors, whereas genetic ablation or silencing suppresses these effects and disrupts reward-cue learning^13,17^. Activation of LHA^GABA^ neurons can also elicit gnawing and biting directed toward non-food objects^18,19^, suggesting recruitment of a broader consummatory or oral-motor program rather than strictly food-specific responses. LHA^GABA^ neurons have also been implicated in arousal control, with activation promoting wakefulness^20,21^.

Neuroanatomical studies have shown that the LHA sends widespread projections throughout the forebrain, midbrain, and brainstem^22–24^. Optogenetic studies have begun to define the behavioral functions of distinct LHA^GABA^ pathways. For example, stimulation of LHA^GABA^ projections to the ventral tegmental area (VTA)^25,26^, paraventricular hypothalamic nucleus (PVH)^27^, diagonal band of Broca (DBB)^28^, and dorsal pons (DP)^29^ promotes feeding. Stimulation of LHA^GABA^→VTA projections also increase gnawing of non-food objects^25^. In addition, activation of LHA^GABA^ projections to the periaqueductal gray (PAG) promotes predatory attack^30,31^. Reinforcement-related effects have likewise been reported following stimulation of LHA^GABA^ projections to the VTA and DBB^26,28^. Together, these studies indicate that distinct LHA^GABA^ pathways can support overlapping functions, particularly in feeding and motivated behavior. However, because most terminal-stimulation studies have examined a limited set of behaviors, they do not yet provide an integrated view of how LHA^GABA^ outputs coordinate motivated behavioral responses. Defining this functional organization requires systematic comparisons of specific projection-targeted stimulation across a broad set of behavior assays to determine whether LHA^GABA^ efferents make distinct contributions or converge on a shared behavioral repertoire.

The DP and lateral preoptic area (LPO) are ideal targets for this comparison because they are prominent, anatomically distant hindbrain and forebrain LHA projection fields, yet their broader behavioral contributions beyond feeding and arousal remain incompletely defined^20,22,29,32^. Here, we first performed histological analyses that found extensive LHA^GABA^ innervation throughout the LPO and DP, with DP fibers distributed across the peri-locus coeruleus, laterodorsal tegmental nucleus (LDT), and Barrington’s nucleus (Bar). We next used optogenetics to systematically compare the effects of activating LHA^GABA^ cell bodies and their projections to the DP and LPO across diverse assays of feeding, non-food object interaction, predatory behavior, real-time place preference, and operant self-stimulation. Across these assays, stimulation of LHA^GABA^ somata and of LHA^GABA^**→**DP and LHA^GABA^**→**LPO terminal fields produced largely overlapping effects, including hyperphagia, gnawing and shredding of non-food objects, killing and consumption of crickets, and reinforcing effects. Together, these findings suggest that major ascending and descending LHA^GABA^ outputs coordinate a shared motivational repertoire across behavioral contexts, rather than supporting strictly segregated pathway-distinct functions.

## MATERIALS AND METHODS

### Ethics Statement

All protocols were performed with the approval of the Institute of Animal Care and Use Committee at the University of Connecticut in accordance with the ethical guidelines described in the National Institutes of Health *Guide for the Care and Use of Laboratory Animals*.

### Animals

Adult male and female mice (2–8 months old) were used in all experiments. *Slc32a1^ires-Cre^* knock-in mice^33^ (The Jackson Laboratory; Stock # 028862) were maintained as heterozygotes on a C57BL/6J background. Mice were housed on a reversed 12 h light/dark cycle (lights off at 9:00 A.M.), and all experiments were conducted during the dark phase. Food and water were available *ad libitum*. Mice were group-housed prior to surgery.

### Surgery

All survival surgeries were performed under 1-3% isoflurane anesthesia using aseptic technique. Mice were placed in a stereotaxic frame (Model 942, Kopf Instruments) on a warming pad (RT-0514, Kent Scientific) to maintain body temperature. Anesthesia was delivered via a nose cone and monitored throughout the procedure. Ketoprofen (5 mg/kg i.p., Covetrus) was administered intraoperatively and at 24- and 48-hours postoperatively for analgesia. For viral delivery, a midline scalp incision was made, the skull was exposed and leveled, and bilateral craniotomies were drilled (OmniDrill 35, WPI). Viral constructs (titers reported as genome copies per milliliter) were loaded into glass micropipettes (Part 21-171, Fisher Scientific), backfilled with mineral oil (Part O1211, Fisher Scientific), and injected using a Nanoliter 2020 (WPI). For optogenetic activation and tracing, *Slc32a1^ires-Cre^* mice received bilateral LHA injections of AAVs expressing Cre-dependent channelrhodopsin-2 (ChR2) with enhanced yellow fluorescent protein (eYFP) (AAV-Ef1α-DIO-ChR2(H134R)-eYFP; serotype 2; 1 × 10¹³ vg/mL; Lot AV4378; UNC Viral Vector Core; *n* = 46 mice) or an eYFP control (AAV-Ef1α-DIO-eYFP; serotype 2; 1.2 × 10¹³ vg/mL; Lot# AV4842, *n* = 31 mice). A volume of 30 nL was delivered at 1 nL/s, with coordinates from bregma as follows: anterior-posterior (AP) −1.50 mm, medial-lateral (ML) ±1.2 mm, and dorsal-ventral (DV) −5.3 mm. Four weeks later, a second surgery was performed for optical implantation. Fiber-optic probes (200 μm, 0.39 NA) were constructed from multimode fiber (FT200EMT, Thorlabs) and housed in ceramic ferrules (MM-CON2007-2300, Precision Fiber Products). Bilateral optical fibers were implanted above the LHA (AP −1.5, ML ±2.0, DV −4.4 mm, 10°), DP (AP −5.4, ML ±1.4, DV −3.0 mm, 10°), or LPO (AP +0.3, ML ±2.6, DV −4.4 mm, 20°), and secured to the skull with Metabond (Parts S371, S398, and S396, Parkell). Mice recovered for at least 1 week prior to behavioral testing.

### Optogenetic stimulation

The optical implants of *Slc32a1^ires-Cre^* mice were connected for photo-stimulation delivery to LHA neurons or their projections to DP or LPO using previously described procedures^34^. Implants were connected via mating sleeves (Thorlabs, ADAL4-5) to patch cables (0.37 NA; 60 cm length; Doric Lenses, Canada, SBP(2)_200/220/900-0.37_0.6_FCM-2xMF1.25) coupled to a fiberoptic rotary joint (Doric Lenses, FRJ 1x1 FC-FC). During behavioral experiments, photo-stimulation (473 nm, 10 ms pulse width, ∼10 mW at the fiber tip) was delivered using a DPSS laser (Shanghai Laser & Optics Century Co, Ltd., China, BL473T8-200FC). Behavior was recorded using a monochrome GigE camera (Noldus) and synchronized with photo-stimulation via a TTL pulse generator (Doric Lenses, OPTG-4; Master-9, MicroProbes) controlled by EthoVision XT v17 software (Noldus). Mice were habituated to tethering at least twice for 20 min each prior to testing.

### Food preference test (FPT)

Mice were habituated in their home cages over two days to grain-based rodent food tablets (20 mg, Test Diets, 5-TUM) or calorie-free cellulose tablets (20 mg, Test Diets, 5-AKU). *Ad libitum* fed mice were placed in a clear-walled PVC arena with grey floor (30 x 30 x 33 cm) for 30 min. Two plastic dishes (5 cm inner diameter) were placed in opposite corners, containing 3 g each of chow or cellulose tablets. The side paired with food was counterbalanced across photo-stimulation sessions (0, 5, 10, and 20 Hz) using a within-subject design. Intake of both food and cellulose was measured by an experimenter blinded to treatment.

### Non-food behavior (NFB)

Mice were habituated in their home cages over two days to four distinct non-food objects: cellulose tablets, a cotton nestlet (Ancare), wood sticks (3 cm; Global Industrial, B776885), and manzanita sticks (3-inch, Bio-Serv, W0016) across 4 days (0–20 Hz). Behavioral testing was conducted under four photo-stimulation sessions (0, 5, 10, and 20 Hz). For testing, mice were placed in a clear-walled PVC arena with a gray floor (30 x 30 x 33 cm) for 30 min. Four plastic dishes were placed in the corners containing 3 g of cellulose pellets, one nestlet, five wooden sticks (3 cm each), and one manzanita stick. Object positions were counterbalanced across photo-stimulation sessions. The total time spent engaging in gnawing (defined as sustained object-directed contact and biting with the incisors; e.g., cellulose, wood, or manzanita) and shredding (defined as tearing or pulling apart nestlet material with the incisors and/or paws) was manually scored by an experimenter blinded to treatment and combined into a single measure. To ensure scoring accuracy and eliminate transient artifacts, a minimum continuous duration of 1 s was required for an active behavior to be scored as a valid bout, whereas inactivity required a minimum of 2 s. Other measured behaviors included total time spent carrying any object (defined as lifting and transporting an item from its original location using the mouth or forelimbs), as well as walking (defined as forward locomotion resulting in whole-body displacement), rearing (defined as raising onto hindlimbs with forelimbs off the floor or against the wall), inactivity (defined as remaining stationary without locomotion or object interaction, excluding grooming), and grooming (defined as self-directed cleaning behaviors such as licking or scratching).

### Cricket hunting (CH)

*Ad libitum* fed, cricket-naïve mice were exposed to ten live house crickets (*Acheta domesticus*) (Petco) in a clear-walled, plexiglass open-field arena (43 x 43 x 33 cm), as previously described^30^. A single 21 min optogenetic session used a block paradigm consisting of ten 1-min light OFF periods interleaved with eleven 1-min ON periods (5, 10, 20 Hz). The number of crickets killed and consumed was measured by an experimenter blinded to treatment condition.

### Real-time place preference (RTPP)

Mice were placed in a two-compartment grey PVC arena (60 x 30 x 33 cm) for 30 min, as previously described^34^. Photo-stimulation was triggered upon entry into one compartment and terminated upon exit. The stimulation-paired compartment was randomly assigned and counterbalanced across sessions (0, 5, and 10 Hz). The percentage of time spent in the stimulation-paired compartment was recorded by EthoVision v17.

### Self-stimulation (SS)

Mice were trained in an operant chamber (23 x 20 x 13 cm) (Med Associates) to lever press for photo-stimulation (3-s, 5 or 10 Hz optical pulse train) in 30-min fixed-ratio (FR1) sessions, as similarly described^13^. For testing, mice were run in the same paradigm to assess optical self-stimulation. Stimulation frequencies (0, 5, or 10 Hz) were tested in a within-subject design. Each 30-min session was initiated by the animal’s first active lever press. Under a FR1 schedule, each lever press delivered a 3-sec optical pulse train during which additional presses were recorded but did not further trigger stimulation. The number of stimulations earned were recorded by MedPC 5 (Med Associates).

### Tissue collection and processing

At the conclusion of the experiments, brain tissue was collected to verify viral expression and optical implant placement. For Fos analysis, a subset of mice received photo-stimulation (10-Hz, 10-ms pulse width) for 30 min and were perfused ∼90 min after photo-stimulation onset. Mice were deeply anesthetized with isoflurane and transcardially perfused with 0.1 M phosphate-buffered saline (PBS), followed by 4% paraformaldehyde in PBS (PFA/PBS). Brains were post-fixed overnight in 4% PFA/PBS at 4°C, cryoprotected in 30% sucrose in PBS for 48 h, embedded in O.C.T. compound (Thermo Fisher), frozen, and stored at −80°C. Coronal sections (40 µm) were cut on a cryostat (Leica CM 3050S) and stored at −20°C in cryoprotectant solution until further processing.

### Immunohistochemistry

Sections were rinsed in 0.1 M PBS and blocked for 2 h at room temperature in 2% normal donkey serum (NDS, Jackson ImmunoResearch).

To visualize neuronal populations of interest for anatomical experiments, sections were incubated overnight at room temperature in blocking solution with primary antibodies, including mouse anti-tyrosine hydroxylase (TH) (1:1000, R&D Systems, Cat# MAB7566), sheep anti-forkhead box protein P2 (FOXP2) (1:1000, R&D Systems, Cat# AF5647), and goat anti-choline acetyltransferase (ChAT) (1:1000; Millipore, Cat# AB144P). Sections were then washed in 0.1% Triton X-100 in PBS and incubated for 2 h at room temperature with secondary antibodies (1:500; Abcam), including donkey anti-mouse Alexa Fluor 405 (Cat #ab175658), donkey anti-sheep Alexa Fluor 594 (Cat # ab150180), and donkey anti-goat Alexa Fluor 594 (Cat # ab150132).

In Fos experiments, eYFP expression in LHA somata was amplified using rabbit anti-green fluorescent protein (GFP) (1:1000, Thermofisher Scientific, Cat# A11122) and donkey anti-rabbit secondary Alexa Fluor 488 (1:500; Abcam, Cat# ab150073). Fos was detected using chicken anti-Fos primary antibody (1:2000; Synaptic Systems, Cat # 226-009) and goat anti-chicken Alexa Fluor 594 secondary (1:500; Abcam, Cat# ab150172).

For both staining protocols, sections were washed after secondary antibody incubation. After final PBS washes, sections were mounted and coverslipped using Vectashield HardSet Antifade Mounting Medium with 4′,6-diamidino-2-phenylindole (DAPI) (Vector Laboratories).

### Digital image processing

To verify viral expression and map projections, immunofluorescent sections were imaged at 10x and 20x magnification using a Keyence epifluorescence microscope (BZ-X800). To assess the spatial relationship between eYFP-positive fibers and immunolabeled neurons, high-magnification images were acquired at 40x with 1-µm z-steps using a Leica SP8 confocal microscope.

For Fos experiments, images were acquired at 20x with 0.28 µm z-steps using a Leica DM6 epifluorescence microscope. Images were collected from the LHA, approximately –1.4 to –1.7 mm relative to bregma, using the fornix as an anatomical landmark and guided by the Paxinos and Franklin mouse brain atlas^35^. To improve image clarity and reduce out-of-focus blur, DM6 images were deconvolved using the Leica Instant Computational Clearing algorithm in LAS X 3.8.1.

Digital images were exported from Keyence (1.3.1.1) or LASX software (v3.7.5) as uncompressed files (.lif or .tif) and processed in Adobe Photoshop 13.0. Z-stacks were converted to maximum-intensity projections. For display only, representative images were adjusted uniformly for brightness and contrast to enhance visualization; unadjusted images were used for quantification.

### Cell counts

Fos quantification was performed on every third 40-μm coronal section through the rostrocaudal extent of the LHA from *Slc32a1^ires-Cre^*mice that received bilateral LHA injections of AAVs expressing Cre-dependent ChR2-eYFP or eYFP and bilateral photo-stimulation via optical fibers targeting the LHA, DP, or LPO. Depending on tissue availability and staining quality, four to six sections were analyzed per subject. LHA boundaries were defined using eYFP immunoreactivity and anatomical landmarks. Images containing Fos+ and DAPI+ fluorescent somata were separated into red and blue channels. For quantification, the middle 30 optical sections from each z-stack of approximately 60 sections were selected. These sections were acquired at 0.28-μm z-steps, corresponding to an 8.4-μm segment centered within the z-stack, and were converted into maximum-intensity projections (MIPs) using a custom FIJI script (v2.0).

Unadjusted MIPs were segmented in Cellpose^36^ (v3.0.10) using a custom-trained deep-learning model. An estimated object diameter of ∼30.8 pixels, corresponding to the approximate soma size at 20x magnification, was used for segmentation based on visual inspection of DAPI+ and Fos+ labeled somata. Segmentation masks were manually verified against the original fluorescence images by an experimenter blinded to experimental group assignment to confirm that selected objects corresponded to individual somata. Fos+ somata were defined as objects containing blue signal within the segmented mask and exhibiting round nuclear labeling consistent with Fos immunoreactivity. Masks were deselected if they corresponded to merged cell bodies, autofluorescent puncta, tissue debris, or non-nuclear Fos-channel objects. Any undetected Fos somata were manually added by an experimenter blinded to group assignment using the aforementioned criteria. Verified masks were then filtered in ImageJ to exclude objects smaller than 113 pixels^2^. Fos+ and DAPI+ cells were quantified from the resulting verified, filtered Cellpose masks. Segmentation and quantification parameters were applied consistently across all images using the same blinded workflow. As a positive control to confirm robust detection of stimulation-induced Fos expression, tissue from mice receiving LHA^GABA^ soma photo-stimulation was included in each immunohistochemistry assay and compared between ChR2 and eYFP groups.

Cellpose outputs were used to calculate the percentage of Fos+/DAPI+ double-labeled somata in each section. Values were then averaged across the rostrocaudal LHA sections for each subject, with the subject used as the unit of analysis. The percentage of double-labeled cells was calculated as: % Fos+/DAPI+ cells = (Fos+/DAPI+ somata / DAPI+ somata) x 100.

### Statistical analysis

Behavioral data were analyzed using two-way repeated measures ANOVAs with Geisser–Greenhouse correction to assess group (ChR2 vs. eYFP) x frequency (0–20 Hz) effects. Separate two-way repeated-measures ANOVAs were used to evaluate implant region (LHA, DP, LPO) x frequency (0–20 Hz) effects in ChR2-expressing mice. Bonferroni *post hoc* tests were applied as appropriate. For Fos analysis, ChR2 and eYFP groups were compared using unpaired *t*-tests separately for each optical-fiber target (LHA, DP, or LPO) following 10-Hz photo-stimulation. Statistical significance was set at *P* < 0.05 (two-tailed). Data are expressed as mean ± SEM. All analyses were conducted in GraphPad Prism 10 or 11, and figures were compiled in Adobe Illustrator 2026.

Exact statistics and sample sizes are listed in the figure legends and Supplemental Material. Data from the following ChR2-expressing mice were excluded from all analyses due to poor viral expression or mistargeted fiber placement: *n*=1 LHA^GABA^, *n*=2 LHA^GABA^→DP, and *n*=2 LHA^GABA^→LPO. Attrition before later assays reduced sample sizes for some groups, including before the real-time place preference test (*n*=2 LHA^GABA^ ChR2; *n*=1 LHA^GABA^→LPO eYFP) and before the cricket hunting assay (n=1 LHA^GABA^ ChR2; n=1 LHA^GABA^→DP eYFP; *n*=1 LHA^GABA^→LPO eYFP). One mouse (LHA^GABA^→LPO ChR2) was excluded only from the self-stimulation analysis because it failed to acquire the operant response. Final sample sizes are reported in the figure legends.

## RESULTS

### Neuroanatomical characterization of LHA^GABA^ projections and optical fiber placements

Prior studies have identified the DP and ventrolateral preoptic area (VLPO) as major projection fields of LHA^GABA^ neurons, implicating these pathways in feeding^29^ and arousal^20,32^. However, the organization of LHA^GABA^ projections across the broader DP and LPO projection fields remains incompletely defined, particularly with respect to rostrocaudal distribution and subregion-specific innervation. To address this gap, we injected a Cre-dependent AAV expressing channelrhodopsin-2 (ChR2)-eYFP into the LHA of *Slc32a1^ires-Cre^*mice^33^ and performed histological analysis of eYFP expression (Fig. 1A–B). Across serial parasagittal and coronal sections, eYFP+ somata were largely restricted to the LHA across its mediolateral and rostrocaudal extent (Fig. 1B–C). We observed a prominent descending projection of LHA^GABA^ neurons through the midbrain, terminating in a dense plexus within the DP (Fig. 1D), consistent with previous work LHA projection patterns^22,24^, and LHA^GABA^ neurons in particular^20^. To resolve innervation patterns within this heterogeneous region^37^, we combined eYFP immunolabeling with markers for noradrenergic neurons (tyrosine hydroxylase; TH), cholinergic neurons (choline acetyltransferase; ChAT), and regional cytoarchitecture (FOXP2). This analysis revealed distinct patterns of LHA^GABA^ innervation across DP subregions. In the rostral mediodorsal DP, corresponding to the LDT, dense varicose eYFP-positive fibers coursed among ChAT-positive cholinergic neurons (Fig. S1A). At the level of the locus coeruleus (LC), LHA^GABA^ terminals were concentrated medial to the LC core, within the peri-LC region, whereas only sparse fibers entered the LC proper among TH-positive noradrenergic neurons (Fig. 1D and Fig. S1B). Using FOXP2 to delineate Bar as a FOXP2-negative zone bordered by FOXP2-positive cells^38^, we identified a dense terminal field ventromedial to the LC core within the Bar region (Fig. 1D and Fig. S1B), consistent with previous work^20^. We also observed extensive ascending LHA^GABA^ projections throughout the rostrocaudal extent of the LPO, extending beyond classical VLPO boundaries (Fig. 1E). Together, these findings refine anatomical descriptions of LHA^GABA^ projections to the LPO and DP, extending preoptic mapping beyond the VLPO and resolving distinct patterns of dorsal pontine innervation.

**Figure 1.**
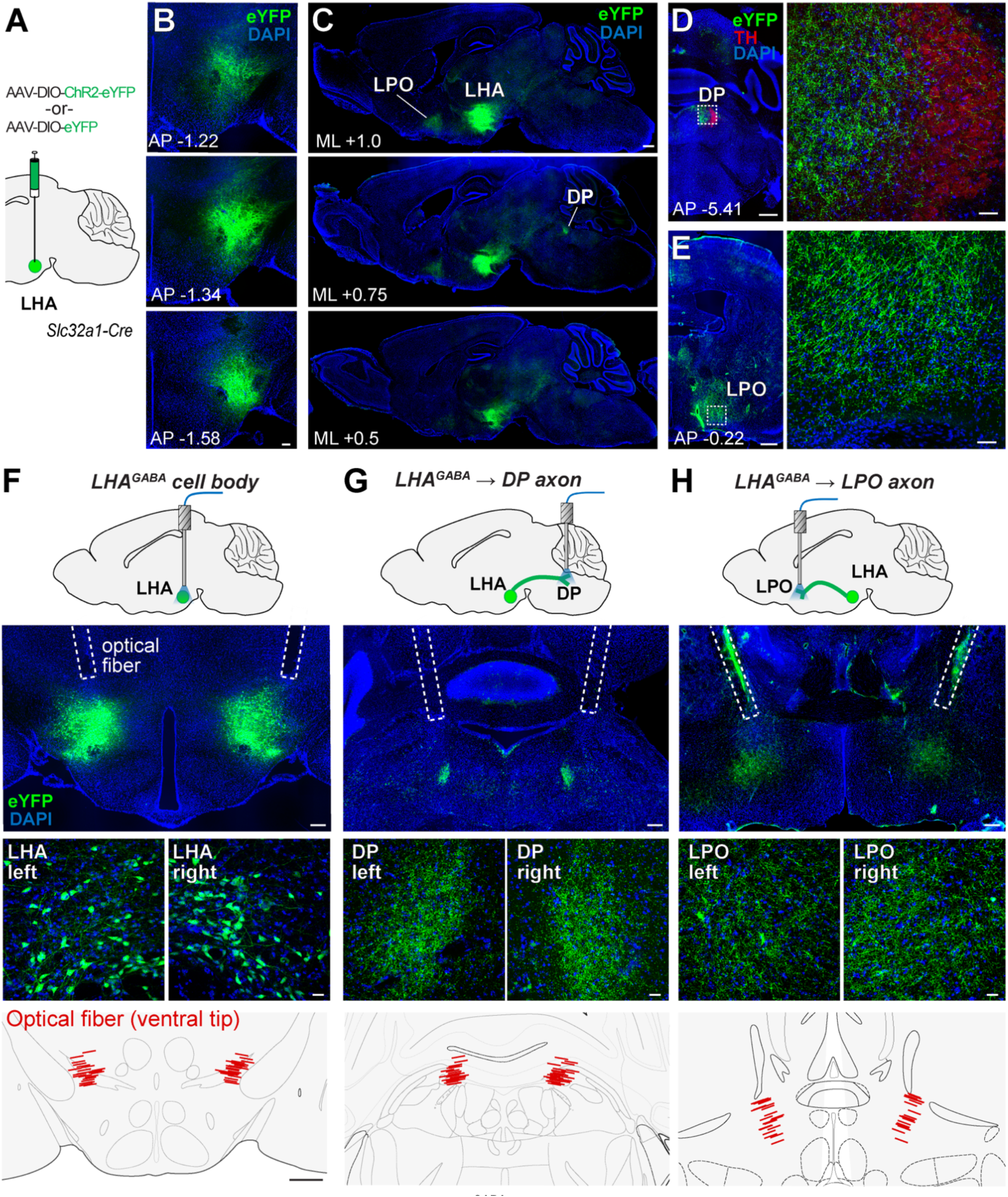
Viral labeling and optogenetic targeting of LHA^GABA^ neurons and their descending and ascending projections to the DP and LPO. **A** Schematic of the viral strategy for Cre-dependent expression of ChR2-eYFP or eYFP in LHA GABAergic neurons of *Slc32a1*-Cre mice. **B** Serial coronal sections, arranged from rostral to caudal (distance from bregma indicated in mm), showing confined eYFP expression in the LHA. Scale bar, 200 µm. **C** Serial parasagittal sections, arranged from lateral to medial (distance from midline indicated in mm) showing brain-wide LHA^GABA^ projections, including prominent terminal fields in the DP and LPO. Scale bar, 500 µm. **D** LHA^GABA^ fibers in the DP relative to tyrosine hydroxylase-positive neurons (TH), shown at low (left) and high magnification (right). Dashed line shows the location of high-magnification imaging. Scale bars, 500 and 25 µm. **E** LHA^GABA^ fibers in the LPO at low (left) and high magnification (right). Dashed line shows the location of high-magnification imaging. Scale bars, 500 and 50 µm. **F** Optogenetic targeting of LHA^GABA^ cell bodies: schematic of bilateral optogenetic fiber placement (row 1), histological verification of expression and fiber tracks at low magnification (row 2) and high magnification (row 3), and summary of fiber tip locations across mice (row 4). Dashed white lines in row 1 show the location of optical fibers in representative mice. Red lines in row 4 show the location of the ventral tips of the optical fibers across subjects. Scale bars, 200, 20, and 500 µm. **G** Same as in (F), for optogenetic targeting of LHA^GABA^ axon terminals in the DP. (H) Same as in (F), for optogenetic targeting of LHA^GABA^ axon terminals in the LPO. AP, anteroposterior; ML, mediolateral.

For optogenetic activation of LHA^GABA^ somata and their projections to the DP and LPO, we injected Cre-dependent AAVs expressing ChR2 or eYFP into the LHA of *Slc32a1^ires-Cre^* mice^33^ and implanted bilateral optical fibers targeting the LHA, DP, or LPO (Fig. 1F–H). These experimental and control mice were used for all subsequent behavioral analysis. Histological validation later confirmed both the injection site and optical fiber placement (Fig. 1F-H).

### Effects of optogenetic activation of LHA^GABA^ neurons and their projections to the DP and LPO on food and cellulose intake

Previous work has shown that activation of LHA^GABA^ neurons^13,39,40^ and their projections to the DP^29^ is sufficient to drive hyperphagia in sated mice. Here, we sought to replicate these established effects and test whether the LHA^GABA^→LPO pathway similarly promotes hyperphagia. Using somatic- and projection-targeted optogenetic stimulation (Fig. 1F–H), we asked whether stimulation selectively increases caloric food intake rather than nonspecific pellet-directed chewing. *Ad libitum*-fed mice were therefore given access to either grain-based food tablets or noncaloric cellulose pellets matched for size and shape during frequency-dependent photo-stimulation (0, 5, 10, and 20 Hz; 10-ms pulse width) (Fig. 2A).

**Figure 2.**
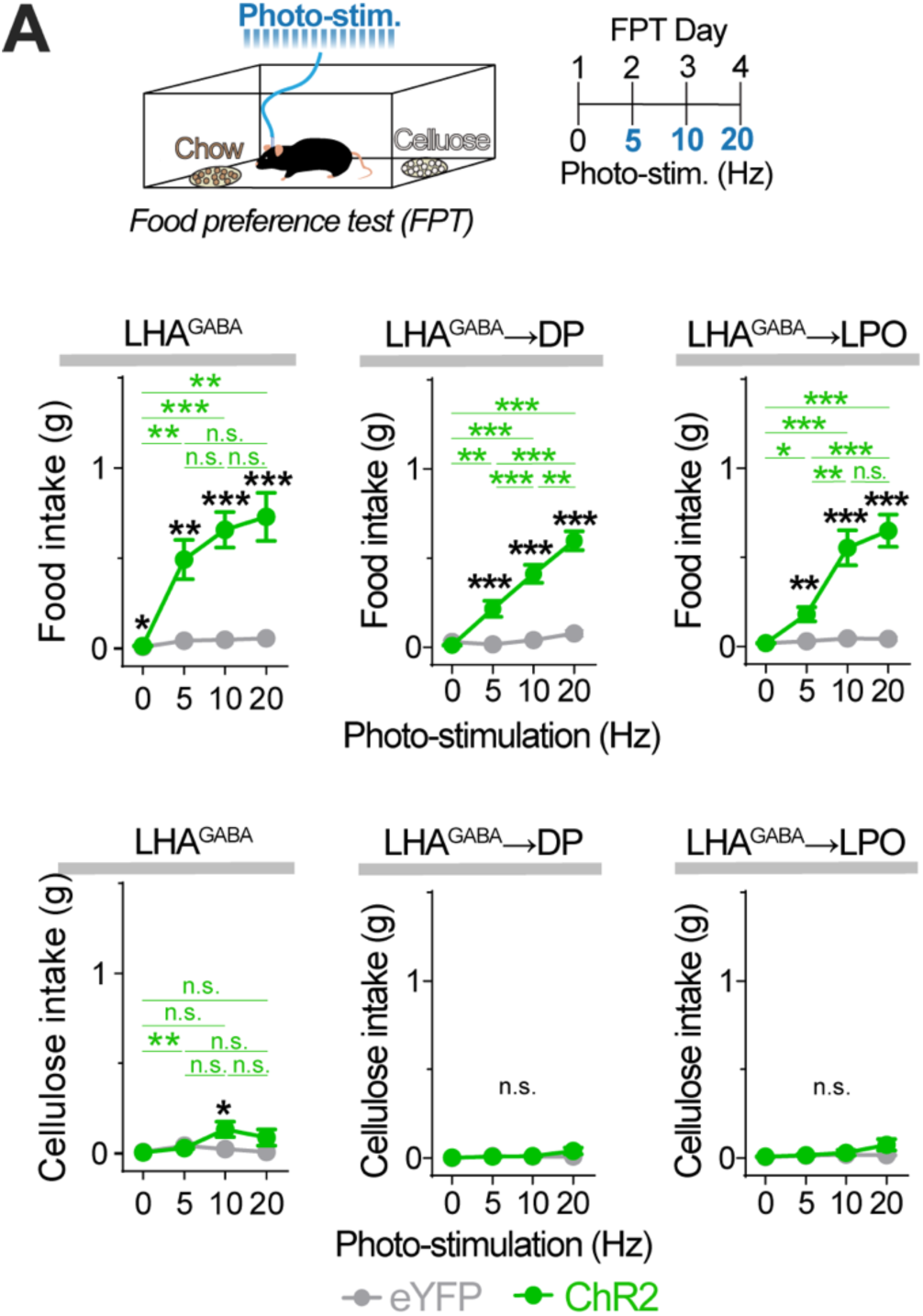
Hyperphagic effects of optogenetic activation of LHA^GABA^ neurons and their projections to DP and LPO. **A** Left. Setup for food preference test (FPT) during photo-stimulation. Right: Experimental timeline. **B** Food intake. Two-way RM ANOVA revealed significant frequency x opsin interaction for LHA^GABA^ (*F*_1.830,38.43_=10.57, *P*=0.0003), LHA^GABA^ → DP (*F*_2.649,58.27_=28.87, *P*<0.0001), and LHA^GABA^ → LPO (*F*_2.245,51.65_=17.80, *P*<0.0001). **C** Cellulose intake. Two-way RM ANOVA revealed a significant frequency x opsin interaction for LHA^GABA^ (*F*_1.961_, _41.18_=4.170, *P*=0.0231), but not for LHA^GABA^ → DP (*F*_1.187, 26.11_=1.444, *P*=0.2457) or LHA^GABA^→LPO (*F*_1.376, 31.64_=1.839, *P*=0.1834). Bonferroni post hoc: \*\*\**P*<0.001, \*\**P*<0.01, \**P*<0.05. Black asterisks indicate ChR2 vs. eYFP comparisons; green asterisks indicate frequency-dependent effects within ChR2 mice; n.s., non-significant. Data are mean ± SEM. Sample sizes: LHA^GABA^ (eYFP, *n*=11; ChR2, *n*=12), LHA^GABA^→DP (eYFP, *n*=9; ChR2, *n*=15), LHA^GABA^→LPO (eYFP, *n*=11; ChR2, *n*=14. Full ANOVA results are provided in Supplemental Material. See Fig. S2 for related results.

Baseline food intake at 0 Hz was low overall but differed between eYFP and ChR2 and groups (*P*=0.0466), primarily due to slightly elevated intake in one eYFP subject that did not meet Tukey’s criteria for exclusion as an outlier (Fig. 2B). At active stimulation frequencies, photo-stimulation produced a robust increase in food intake in ChR2-expressing LHA^GABA^, LHA^GABA^→DP, and LHA^GABA^→LPO mice relative to eYFP controls (Fig. 2B). In all three ChR2 groups, food intake increased with stimulation frequency (Fig. 2B).

Cellulose intake did not differ between ChR2 and eYFP groups at 0 Hz (Fig. 2C). At active stimulation frequencies, cellulose intake was modestly elevated only in ChR2-expressing LHA^GABA^ mice at 10-Hz, but not at 5 Hz or 20 Hz, and was unchanged in LHA^GABA^→DP and LHA^GABA^→LPO mice at all frequencies (Fig. 2C). Among ChR2-expressing mice, the magnitude of food and cellulose intake did not differ across the LHA^GABA^, LHA^GABA^→DP, and LHA^GABA^→LPO stimulation conditions (Fig. S2). These findings replicate prior evidence that activation of LHA^GABA^ neurons and LHA^GABA^ → DP projections are sufficient to elicit hyperphagia in sated mice and further show that the LHA^GABA^→LPO pathway shares this role. Low cellulose consumption indicates that, when food is available, the hyperphagic response is food-directed rather than a nonspecific increase in ingestive behavior.

### Effects on gnawing, shredding, and other non-food behaviors

Given that activation of LHA^GABA^ neuronal somata, as well as stimulation of the LHA^GABA^→VTA pathway induces gnawing of non-food objects^13,18,25^, we next asked whether stimulation of LHA^GABA^ terminals in the DP and LPO would similarly recruit repetitive gnawing and shredding responses toward non-food objects when food was absent. To address this question, we used a behavioral assay in which mice were allowed to interact with four distinct non-food objects: cellulose pellets, a manzanita stick, wood sticks, and a cotton fiber nestlet (Fig. 3A). The assay was repeated using four photo-stimulation frequencies (0, 5, 10, and 20 Hz). We quantified the time spent gnawing the first three objects, shredding of the nestlet, object carrying, grooming, walking, rearing, and inactivity to capture the broader behavioral repertoire expressed during stimulation.

**Figure 3.**
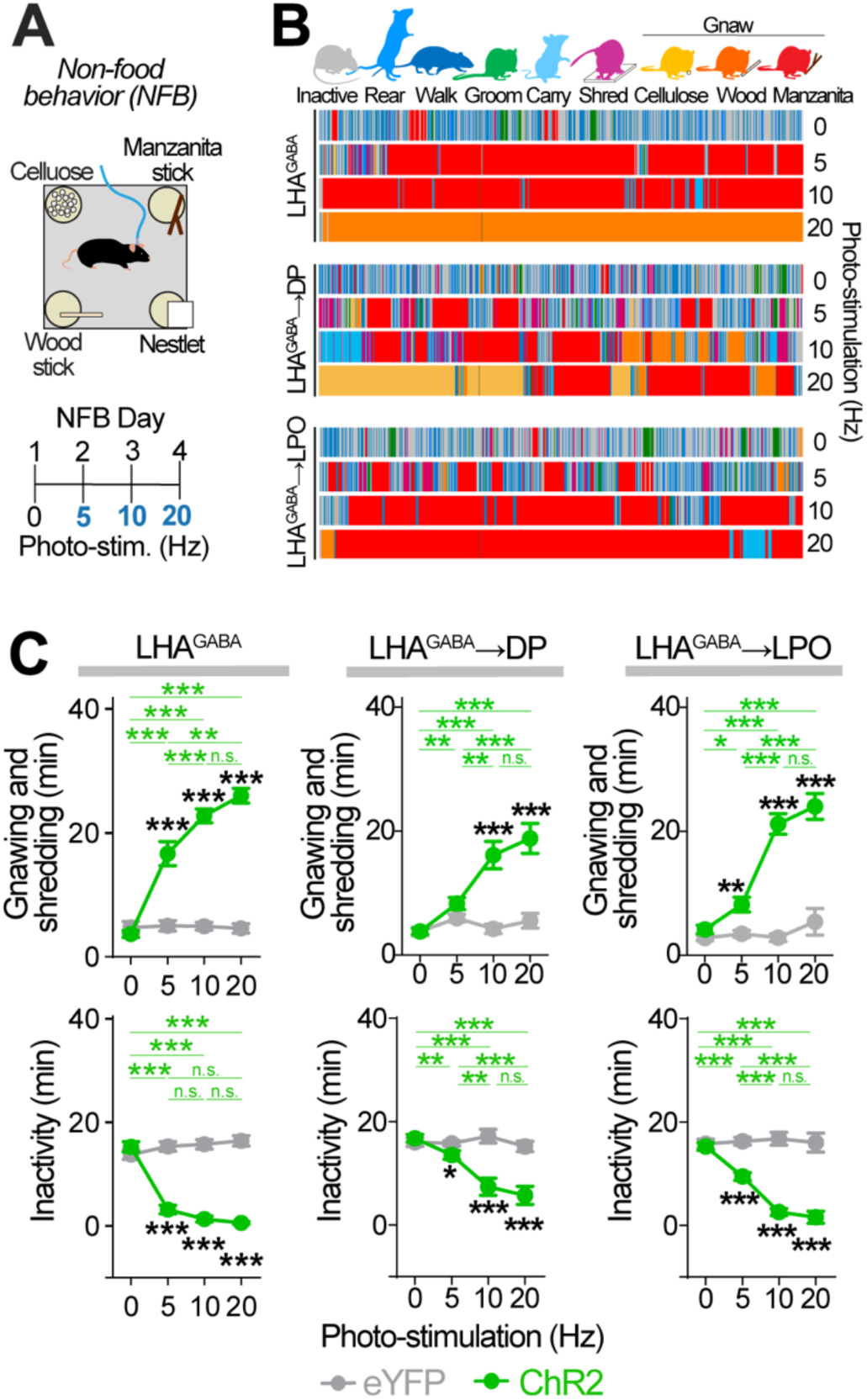
Optogenetic activation of LHA^GABA^ neurons and their projections to DP and LPO increases gnawing and shredding while reducing inactivity. **A** Left. Setup for non-food behavior (NFB) test during photo-stimulation. Right: Experimental timeline. **B** Color-coded behavioral sequences from representative ChR2 mice at different photo-stimulation frequencies. **C** Top: Gnawing and shredding time. Two-way RM ANOVA revealed significant frequency x opsin interactions for LHA^GABA^ (*F*_1.963, 41.22_=56.22, *P*<0.0001), LHA^GABA^→ DP (*F*_1.787, 39.31_=15.36, *P*<0.0001), and LHA^GABA^ → LPO (*F*_1.914, 44.02_=31.33, *P*<0.0001). Bottom: Inactivity time. Two-way RM ANOVA revealed significant frequency x opsin interactions for LHA^GABA^ (*F*_2.294, 48.17_=63.03, *P*<0.0001), LHA^GABA^ → DP (*F*_1.707, 37.55_=16.23, P<0.0001), and LHA^GABA^→LPO (*F*_2.068, 47.56_=33.22, P<0.0001). Bonferroni post hoc: \*\*\**P*<0.001, \*\**P*<0.01, \**P*<0.05. Black asterisks indicate ChR2 vs. eYFP comparisons; green asterisks indicate frequency-dependent effects within ChR2 mice; n.s., non-significant. Data are mean ± SEM. Sample sizes: LHA^GABA^ (eYFP, *n*=11; ChR2, *n*=12), LHA^GABA^→DP (eYFP, *n*=9; ChR2, *n*=15), and LHA^GABA^ → LPO (eYFP, *n*=11; ChR2, *n*=14). Full ANOVA results are provided in *Supplemental Material.*See Fig. S3 and Fig. S4 for releated behavioral results.

Representative time-resolved behavioral ethograms from ChR2-expressing mice showed that, under baseline conditions (0 Hz), behavior consisted primarily of brief bouts of inactivity, rearing, walking, and grooming, with occasional episodes of gnawing, shredding, or carrying non-food objects (Fig. 3B). During photo-stimulation (5, 10, and 20-Hz), behavior shifted toward prolonged bouts of gnawing, interspersed with shorter bouts of shredding and other behaviors (Fig. 3B).

Quantification across animals confirmed that photo-stimulation robustly increased time spent gnawing and shredding while decreasing inactivity in ChR2-expressing LHA^GABA^, LHA^GABA^ → DP, and LHA^GABA^ → LPO mice in comparison to eYFP controls (Fig. 3C). In each group, these interactions increased with stimulation frequency (Fig. 3C). Notably, ChR2-expresing LHA^GABA^ mice spent more time gnawing and shredding than both LHA^GABA^→DP and LHA^GABA^ → LPO mice at 5 Hz, and more than LHA^GABA^→DP mice at 10 and 20 Hz (Fig. S3), suggesting that this behavior is more readily evoked by somatic stimulation than by selective terminal stimulation of either projection.

Modest but significant frequency-dependent reductions in walking and rearing were observed in ChR2-expressing LHA^GABA^, LHA^GABA^→DP, and LHA^GABA^→LPO mice relative to eYFP controls (Fig. S4A–B). A frequency-dependent reduction in grooming behavior was observed only in ChR2-expressing LHA^GABA^→DP mice (*P*<0.05), but not LHA^GABA^ mice (*P*=0.13), or LHA^GABA^→LPO mice (*P*=0.14) (Fig. S4C). Finally, carrying of non-food objects showed a frequency-dependent increase in ChR2-expressing LHA^GABA^, LHA^GABA^→DP, and LHA^GABA^→LPO mice relative to eYFP controls (Fig. S4D). Together, these findings indicate that, in the absence of food, activation of LHA^GABA^ neurons or their projections to the DP or LPO produces non-food-object-directed repetitive motor patterns, including gnawing and shredding, while reducing other behaviors such as walking, rearing, and grooming. These results further show that the LHA^GABA^→LPO and LHA^GABA^→DP pathways drive overlapping behavioral responses.

### Effects on predatory behaviors

LHA^GABA^ neurons can promote complex behavioral sequences, such as predation. Fiber photometry studies have shown that LHA^GABA^ neurons are activated at predatory attack onset and that optogenetic stimulation of these neurons, as well as their projections to the PAG, promotes predation- related behavior^30,31^. We therefore asked whether stimulation of LHA^GABA^ terminals in the DP and LPO is likewise sufficient to enhance predatory behavior. To test this, *ad libitum*-fed ChR2- and eYFP-expressing mice were given access to live crickets during photo-stimulation (5, 10, and 20 Hz) in a cricket hunting assay (Fig. 4A). Photo-stimulation increased the number of crickets killed and consumed at each frequency in ChR2-expressing LHA^GABA^, LHA^GABA^ → DP, and LHA^GABA^ → LPO mice relative to eYFP controls (Fig. 4B). Within ChR2 groups, predatory behavior increased across stimulation frequencies (Fig. 4B). However, eYFP-expressing controls also showed increased predatory behavior across testing days, particularly in the LHA^GABA^ cohort, in which significant differences were observed between 5-Hz and 20-Hz sessions (Fig. 4B). These within-group differences in eYFP controls are likely attributable to increased hunting experience, as the three stimulation frequencies were tested on consecutive days in a fixed order (5, 10, and 20 Hz). Therefore, the apparent frequency-related increase in predation in the ChR2 group may reflect both photo-stimulation and repeated hunting experience and should be interpreted with caution. Direct comparisons among ChR2 groups showed that LHA^GABA^ mice consumed more crickets than LHA^GABA^→DP and LHA^GABA^→LPO mice at 10 Hz (Fig. S5), suggesting that consummatory aspects of predatory behavior may be more readily evoked by somatic stimulation than by selective terminal stimulation of either projection. Together, these data indicate that both LHA^GABA^→DP and LHA^GABA^→LPO projections are sufficient to enhance predatory behavior.

**Figure 4.**
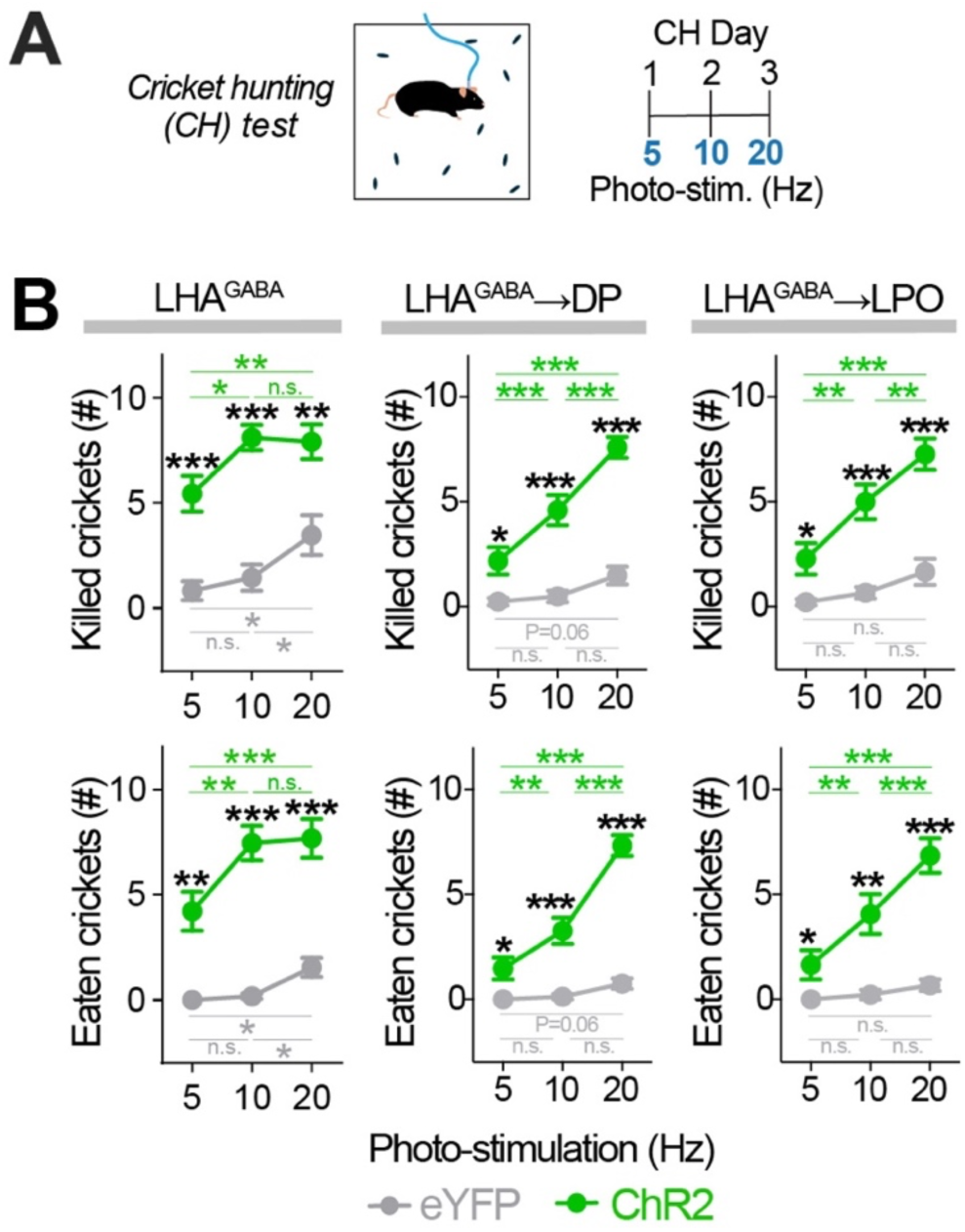
Optogenetic activation of LHA^GABA^ neurons and their projections to DP and LPO increases hunting behavior. **A** Left: Setup for the cricket hunting (CH) test during photo-stimulation. Right: Experimental timeline. **B** Top: Number of crickets killed. Two-way RM ANOVA revealed a trend toward a frequency x opsin interaction in LHA^GABA^ mice (*F*_1.8, 33_=4.0, *P=*0.031) and significant frequency x opsin interactions in the LHA^GABA^ → DP (*F*_1.8, 39_=15.0, *P*<0.001) and LHA^GABA^ → LPO mice (*F*_1.5, 32_=7.9, *P=*0.003). Bottom: Number of crickets eaten. Two-way RM ANOVA revealed significant frequency x opsin interactions in LHA^GABA^ (*F*_1.6, 29_=9.5, *P=*0.001), LHA^GABA^ → DP (*F*_1.8, 38_=25.0, P<0.001), and LHA^GABA^ → LPO mice (*F*_1.8, 38_ =18.0, P<0.001). Bonferroni post hoc: \*\*\**P*<0.001, \*\**P*<0.01, \**P*<0.05. Black asterisks indicate ChR2 vs. eYFP comparisons; green asterisks indicate frequency-dependent effects within ChR2 mice; gray asterisks indicate frequency-dependent effects within eYFP mice; n.s., non-significant. Data are mean ± SEM. Sample sizes: LHA^GABA^ (eYFP, *n*=11; ChR2, *n*=9), LHA^GABA^→DP (eYFP, *n*=8; ChR2, *n*=15), and LHA^GABA^→LPO (eYFP, *n*=9; ChR2, *n*=14). Full ANOVA results are provided in the *Supplemental Material*. See Fig. S5 for related behavioral results.

### Effects on real-time place preference and self-stimulation

As activation of LHA^GABA^ neuronal somata, and particularly their projections to the VTA, has been shown to be reinforcing^13,25,26^, we next asked whether stimulation of LHA^GABA^ terminals in the DP and LPO would similarly support positive valence and reinforcement-related responding in real-time place preference (RTPP) and operant self-stimulation (SS) assays (Fig. 5A–B). In the RTPP assay, photo-stimulation increased time spent in the stimulation-paired zone at each active frequency tested (5 and 10 Hz) in ChR2-expressing LHA^GABA^, LHA^GABA^→DP, and LHA^GABA^→LPO mice relative to eYFP controls (Fig. 5C). Within ChR2 groups, a frequency-dependent increase in time spent in the stimulation-paired zone was observed (Fig. 5C). Direct comparisons among ChR2 groups showed that LHA^GABA^ mice spent more time in the stimulation-paired zone than both LHA^GABA^→DP and LHA^GABA^→LPO mice at 5 Hz, and more than LHA^GABA^→DP mice at 10 Hz (Fig. S6), suggesting that RTPP is more readily evoked by somatic stimulation than by selective terminal stimulation of either projection. At 10 Hz, LHA^GABA^→DP mice also spent less time in the stimulation-paired zone than LHA^GABA^→LPO mice (Fig. S6), indicating differences in the effects of projection-targeted stimulation at higher frequencies.

**Figure 5.**
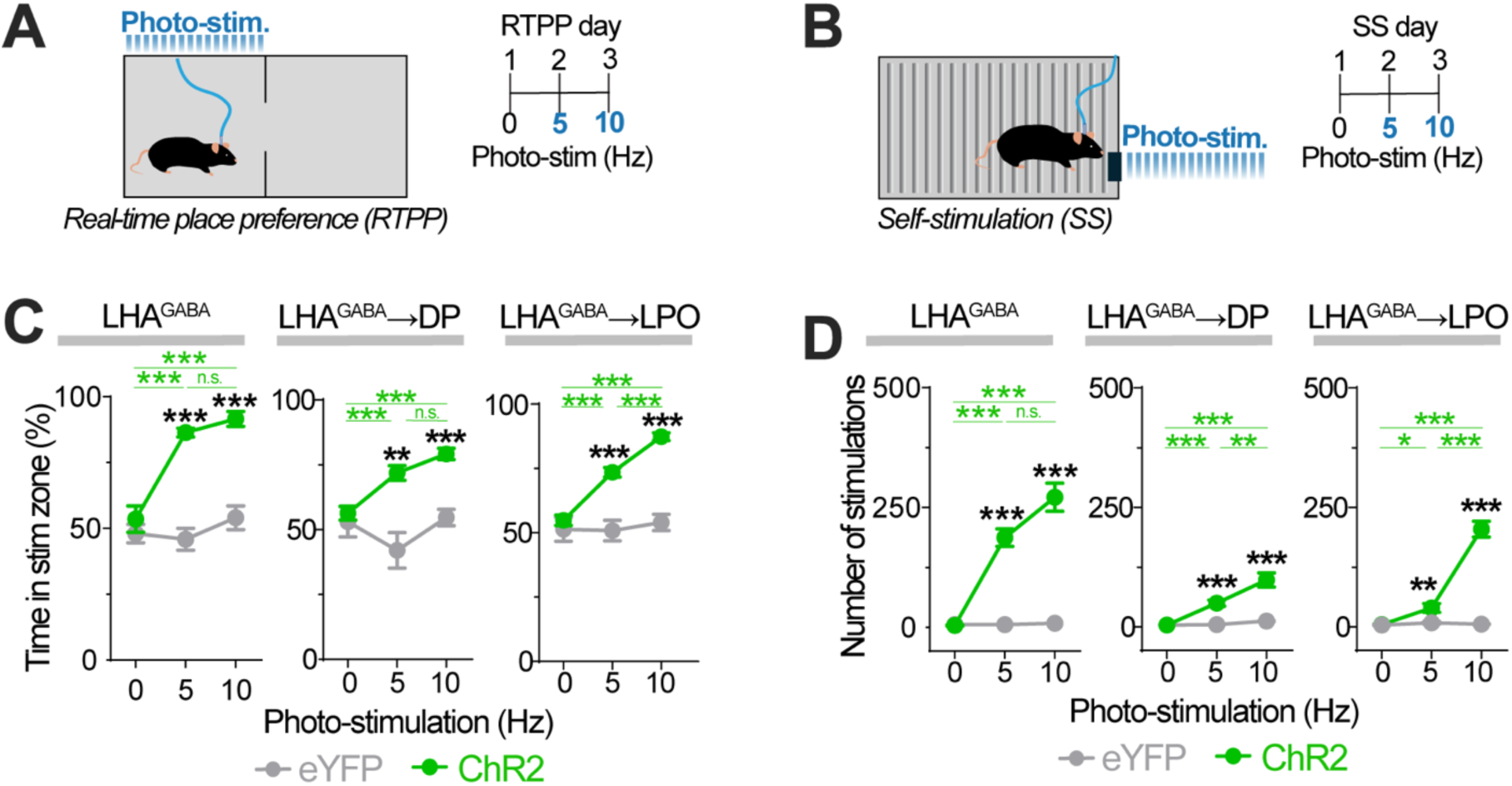
Reinforcing effects of optogenetically activating LHA^GABA^ neurons and their projections to DP and LPO. **A** Left: Setup for the real-time place preference (RTPP) test during photo-stimulation. Right: Experimental timeline. **B** Time in stimulation-paired zone (%). Two-way RM ANOVA revealed significant frequency x opsin interactions for LHA^GABA^ (*F*_1.865, 35.44_=11.62, *P*=0.0002), LHA^GABA^→DP (*F*_1.973, 43.40_=10.23, *P*=0.0002), and LHA^GABA^→LPO (*F*_1.642, 36.13_=16.09, *P*<0.0001). **C** Left: Setup for the self-stimulation test (SS). Right: Experimental timeline. **D** Number of stimulations. Two-way RM ANOVA revealed significant frequency x opsin interactions for LHA^GABA^ (*F*_1.334, 37.36_=54.79, *P*<0.0001), LHA^GABA^ → DP (*F*_1.301, 27.98_=13.69, *P*=0.0004), and LHA^GABA^→LPO (*F*_1.495, 30.65_=71.02, *P*<0.0001). Bonferroni post hoc: \*\*\**P*<0.001, \*\**P*<0.01, \**P*<0.05. Black asterisks indicate ChR2 vs. eYFP comparisons; green asterisks indicate frequency-dependent effects within ChR2 mice; n.s., non-significant. Data are mean ± SEM. Sample sizes—RTPP: LHA^GABA^ (eYFP, *n*=11; ChR2, *n*=10), LHA^GABA^→DP (eYFP, *n*=9; ChR2, *n*=15), LHA^GABA^→LPO (eYFP, *n*=10; ChR2 *n*=14). Sample sizes—SS: LHA^GABA^ (eYFP, *n*=11; ChR2, *n*=10) LHA^GABA^→DP (eYFP, *n*=9; ChR2, *n*=15), LHA^GABA^→LPO (eYFP, *n*=10; ChR2, *n*=13). Full ANOVA results are provided in *Supplemental Material*.

In the SS assay, photo-stimulation likewise increased the number of self-stimulation events at each active frequency tested (5 and 10 Hz) in ChR2-expressing LHA^GABA^, LHA^GABA^→DP, and LHA^GABA^→LPO mice relative to eYFP controls (Fig. 5D). In ChR2-expressing groups, self-stimulation increased with stimulation frequency (Fig. 5D). Direct comparisons among ChR2 groups showed that LHA^GABA^ mice had more self-stimulations events than both projection groups at 5 Hz, and more than LHA^GABA^ → DP mice at 10 Hz (Fig. S6), suggesting that reinforcement-related responding is more readily evoked by somatic stimulation than by selective terminal stimulation of either projection. At 10 Hz, LHA^GABA^ → DP mice also had fewer self-stimulation events than LHA^GABA^→LPO mice (Fig. S6), indicating differences in projection-targeted stimulation at higher frequencies. Together, these findings show that LHA^GABA^ projections to both the DP and LPO support positive valence and are reinforcing.

### Effects on LHA somatic Fos expression

The shared behavioral effects of stimulating LHA^GABA^→DP and LHA^GABA^→LPO terminals may be explained by multiple circuit-level mechanisms. One consideration when interpreting our optogenetic stimulation of projection fields is possible backpropagating activation of LHA somata following terminal stimulation. To determine whether stimulation of LHA^GABA^ terminal fields activates LHA somata, we delivered optogenetic stimulation (10 Hz, 473 nm, 10-ms pulse width) before tissue collection and measured Fos immunoreactivity (IR) in the LHA of a subset of experimental mice. As expected, 10-Hz photo-stimulation increased the percentage of Fos+/DAPI+ double-labeled LHA somata in ChR2-expressing LHA^GABA^ mice compared with eYFP controls (Fig. S7). Following photo-stimulation, ChR2-expressing LHA^GABA^→DP mice had a similar percentage of Fos+/DAPI+ LHA somata as controls (Fig. S7). In contrast, LHA^GABA^→LPO photo-stimulation increased this measure (Fig. S7). These data suggest that 10-Hz stimulation of the LPO, but not DP, terminal-field may recruit LHA somata antidromically, although the underlying mechanism remains uncertain (see *Discussion*).

## DISCUSSION

In this study, we systematically compared optogenetic stimulation of LHA^GABA^ somata with terminal stimulation of two major projection fields, the DP and LPO, across multiple assays of motivated behavior. A principal finding is that activation of LHA^GABA^ somata or their projection fields to the DP and LPO produced broadly overlapping effects across feeding, non-food behavior, predatory behavior, and reinforcement-related assays. Across all three stimulation conditions, activation promoted robust feeding in sated mice and increased oral-motor behaviors such as gnawing and shredding of non-food objects. It also enhanced predatory behavior, as reflected by increased cricket killing and consumption in sated mice. In addition, activation of all three targets supported real-time place preference and operant self-stimulation, indicating that these circuits support positive valence and are reinforcing. Anatomically, LHA^GABA^ fibers extended beyond VLPO boundaries across the broader lateral preoptic field and were distributed across multiple dorsal pontine subregions, including peri-LC, LDT, and Bar. Together, our results extend prior work by demonstrating that stimulation of LHA^GABA^ projections to DP and LPO produce overlapping motivated behaviors across multiple domains. More broadly, these findings may suggest a distributed functional hypothalamic organization in which major ascending and descending LHA^GABA^ pathways engage a shared motivational repertoire rather than wholly distinct behavioral functions.

### LHA as a motivational output system

Our findings build on prior optogenetic or chemogenetic studies showing that stimulation of LHA^GABA^ somata promotes feeding and reinforcement^13^, as well as oral-motor behaviors directed toward non-food stimuli^18^. Projection-specific studies show that multiple LHA^GABA^ outputs promote feeding, including projections to the VTA^25,26^, PVH^27^, DBB^28^, and DP^29^. Increased gnawing of non-food objects has also been observed following stimulation of LHA^GABA^→VTA pathways^25^. Similarly, LHA^GABA^ projections to the PAG promote predatory attack^30,31^, whereas projections to the VTA and DBB support reinforcement-related behaviors^26,28^. Viewed in this context, our findings are notable because stimulation of LHA^GABA^ terminals in two prominent, though divergent, downstream targets—the DP and LPO—produced broadly overlapping effects across feeding, non-food behavior, predatory hunting, and reinforcement-related assays. These results argue against a one-pathway-one-function organization across the projection targets examined here. Instead, they suggest that multiple LHA^GABA^ pathways may contribute to overlapping components of motivated behavior, with the specific behavioral outcome shaped by context and stimulus availability.

### Interpretation of specific features of the behavioral phenotype

Several aspects of the behavioral phenotype provide insight into how LHA^GABA^ circuits may influence behavioral selection. First, the strong preference for caloric food over noncaloric cellulose pellets across somatic and terminal stimulation conditions suggests that these manipulations do not simply produce indiscriminate oral-motor output or nonspecific ingestion. Second, LHA^GABA^ activation appears to shift behavioral selection toward sustained engagement with salient environmental stimuli, even under conditions that would ordinarily constrain those responses, such as satiety or prey novelty^30,31,41^. Third, these same pathways supported real-time place preference and self-stimulation responding, suggesting a potential link between the regulation of innate motivated behaviors and reinforcement learning, consistent with previous work^17,42,43^. Fourth, within an appetitive–consummatory framework^5,7,9,44,45^, the LHA^GABA^ pathways examined here appear to influence multiple phases of motivated action, including consummatory output and appetitive sequences (e.g., prey pursuit and killing). This interpretation is broadly consistent with prior work suggesting that LHA^GABA^ neurons contribute to multiple phases of feeding-related behavior^13,18,25,29–31,46–49^.

### Interpretation of the projection-targeted stimulation

A central question raised by these findings is why activation of LHA^GABA^ projections to the DP and LPO produced such similar behavioral effects. One possibility is that terminal-field stimulation recruited LHA somata through antidromic, or backpropagating activity, particularly following LPO stimulation, thereby activating projections to non-stimulated regions. Supporting this interpretation, our Fos results show that stimulation of the LHA^GABA^→LPO terminal field increased LHA somata activation. This effect was absent following stimulation of the LHA^GABA^→DP terminal field, which may be attributable to many factors, such as the greater anatomical distance between the stimulated terminal field and the LHA. Antidromic activation could result in shared behavioral effects if there are shared influences on, or interactions between, DP and LPO circuitry. First, some LHA^GABA^ neurons may collateralize to both DP and LPO, such that stimulation in one terminal field could recruit neurons that also innervate the other target, or additional downstream targets. Indeed, single LHA hypocretin/orexin neurons have extensive ascending and descending axonal projections^50^, raising the possibility that similar collateralization occurs within LHA^GABA^ neurons. Second, distinct DP- and LPO-projecting LHA^GABA^ neurons could recruit one another through local intra-LHA circuits. Although local synaptic connectivity among LHA neurons has been shown to be exceptionally sparse^51^, other studies support the possibility of broader polysynaptic intra-LHA or regional interactions^52,53^.

The shared behavioral effects could also be explained without invoking antidromic activation, but rather by the functional organization of LHA downstream targets, such as (1) redundancy across downstream targets and/or (2) recruitment of a shared multi-regional network through distinct circuit nodes. Although the present data do not distinguish between these possibilities, both suggest feeding, gnawing/shredding, predation, and reinforcement are regulated by a distributed network that includes the DP and LPO.

Prior studies have implicated the DP and LPO in functions relevant to the motivated behaviors examined here, suggesting that both regions are positioned to influence overlapping behavioral outputs. Consistent with this broader functional role, LHA^GABA^ fibers extended throughout the LPO and VLPO, suggesting potential engagement of LPO functions beyond the well-characterized arousal effects mediated by the VLPO^20,32^.The DP projection field includes several functionally relevant subregions, including the peri-LC region, LDT, and Bar, positioning it to influence arousal/wakefulness^54,55^, reward and motivation^56^, and autonomic/motor output^38,57^. Similarly, the LPO is positioned to influence reward and action selection through its effects on VTA circuitry. Activation of the LPO promotes reward-seeking and modulates VTA activity by inhibiting GABA neurons and disinhibiting dopamine neurons^58^. Both the LPO and its projections to the VTA can support operant reinforcement^59^. Additionally, LPO neurons are engaged during nest-building^60^, a behavior that may be related to the gnawing and shredding observed here.

The strong functional convergence between ascending and descending LHA^GABA^ pathways suggests that their diverse behavioral effects may be supported by fine-grained cellular and projection-level heterogeneity. LHA^GABA^ neurons comprise molecularly distinct subpopulations, including neurons defined by leptin receptor (LepR), neurotensin, galanin, and somatostatin expression^1,15,16,61–65^. Recent work indicates that some of these populations, particularly LHA^LepR^ neurons, can differentially contribute to appetitive and consummatory aspects of feeding^46,47,66^. More broadly, neurotensin- and galanin-expressing LHA^GABA^ subpopulations have been implicated in specialized functions related to energy balance, reward, locomotor activity, and anxiety-like behavior^63,64,67–70^. In the current study, stimulation of each pathway may have engaged multiple LHA^GABA^ subpopulations, contributing to the broad behavioral profile observed here. However, because we did not determine the molecular identities of the activated LHA^GABA^ neurons beyond their GABAergic phenotype or determine whether individual neurons collateralize to both targets, this interpretation remains tentative.

### Limitations of the study

While our Fos analysis suggests possible antidromic recruitment of LHA somata, this approach has important limitations. In this analysis, we quantified the percentage of Fos+/DAPI+ double-labeled LHA somata rather than Fos expression in LHA^GABA^ neurons, because the ChR2-eYFP expression pattern prevented reliable identification of those neurons. Moreover, because Fos is an indirect and threshold-dependent marker of activity, these data cannot establish whether terminal stimulation antidromically activated LHA^GABA^ neurons, only that such recruitment was not uniform across terminal fields and may vary with the distance between the stimulated terminal field and the LHA.

Finally, our experiments were conducted under a limited set of physiological conditions, and LHA^GABA^ function may differ during states such as hunger or stress, which were not directly examined. A related limitation is that the cricket hunting assay did not dissociate prey approach, attack, and consumption at a microstructural level, and frequency-dependent effects in that task were confounded by repeated prey-hunting experience across sessions. As a result, our interpretation of the hunting phenotype is necessarily framed at the level of broad behavioral output rather than fine-grained sequencing.

### Conclusions and future directions

In sum, LHA^GABA^ circuits may engage a broad motivational repertoire rather than strictly promoting discrete behavioral outputs. The overlapping effects observed across ascending and descending projection fields suggests that LHA^GABA^ neurons function within a distributed network, including the DP and LPO, that promotes behavioral engagement across contexts, refining our understanding of hypothalamic output organization.

Several key questions remain. First, determining whether individual LHA^GABA^ neurons collateralize to multiple downstream targets, or instead act through distinct but interacting populations, will require projection-specific tracing and recording approaches. Second, dissecting molecularly defined LHA^GABA^ subpopulations and their projection-specific contributions will be critical for resolving how diverse behavioral functions are coordinated within this system. Third, future work should determine how internal states such as hunger, stress, and learning history modulate the recruitment and function of these pathways. Finally, understanding how these pathways are altered in pathological states may provide insight into maladaptive motivated behaviors, including compulsive feeding and addiction.

## Supporting information

Supplemental Material_Statistics

## ACKNOWLEDGEMENTS

We thank L. Mickelsen, E. Burleigh, S. Manos, E. Beltrami, and N. Taweh for initial work optimizing home-cage behavioral scoring and analysis; M. Kadian for behavioral scoring support; E. Gordon, C. Collier, J. Lesser, and L. Boyd for histology support; C. O’Connell for imaging support; L. Novak for histology and imaging support; B. Dyer for automation of the Fos analysis; C. Kangus for genotyping support. We acknowledge the University of Connecticut Animal Care Services for animal husbandry support, the Advanced Light Microscopy Facility (RRID:SCR_027547) for imaging guidance, and the UConn Mechanical/Glass Facility for custom behavioral boxes.

## AUTHOR CONTRIBUTIONS

Investigation: Y.H. with guidance from W.F., A.C.J. and N.R.S.; formal analysis: Y.H.; data visualization: Y.H., A.C.J., N.R.S.; additional histological analysis: O.K.; writing – original draft: Y.H., A.C.J., N.R.S.; writing - review & editing, Y.H., W.F., A.C.J., N.R.S.; conceptualization, project administration, supervision and funding acquisition: A.C.J. and N.R.S.

## FUNDING

This work was supported by the National Institutes of Health grants R01 DC023564 and R00 DK119586 to N.R.S., R01 MH112739 to A.C.J.; the UConn CLAS Research Equipment Fund and Startup Funds to N.R.S. and NIH Shared Instrumentation Grant S10OD016435 to A. Nishiyama for imaging support.

## COMPETING INTERESTS

The authors declare no competing interests.

## SUPPLEMENTARY INFORMATION

**Figure S1.**
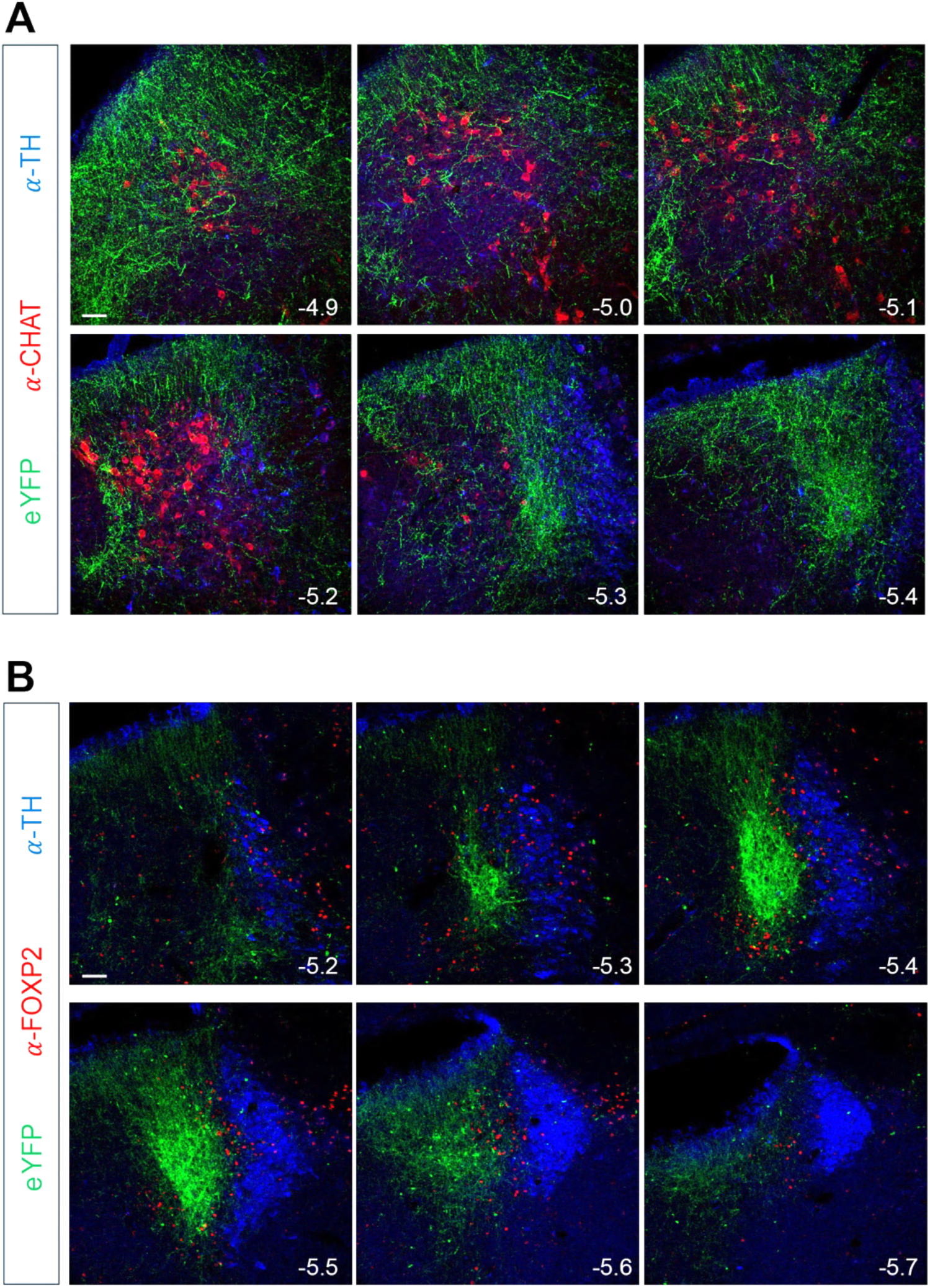
LHA^GABA^ axons densely innervate multiple neuromodulatory nuclei in the dorsal pons. **A** Representative serial coronal sections through the DP, arranged from rostral to caudal (distance from bregma indicated in mm), showing LHA^GABA^ axon terminals (eYFP), ChAT-positive neurons in the LDT, and TH-positive neurons in the LC. Scale bar, 50 µm. **B** Representative serial coronal sections through the DP, from rostral (left) to caudal (right), showing LHA^GABA^ terminals (eYFP), the transcription factor FOXP2, and TH-positive neurons. LHA^GABA^ terminals are densest in a FOXP2-negative zone ventromedial to the TH-positive LC, corresponding to Barrington’s nucleus (Bar). Scale bar, 50 µm. ChAT, choline acetyltransferase; LC, locus coeruleus; LDT, laterodorsal tegmental nucleus; TH, tyrosine hydroxylase.

**Figure S2.**
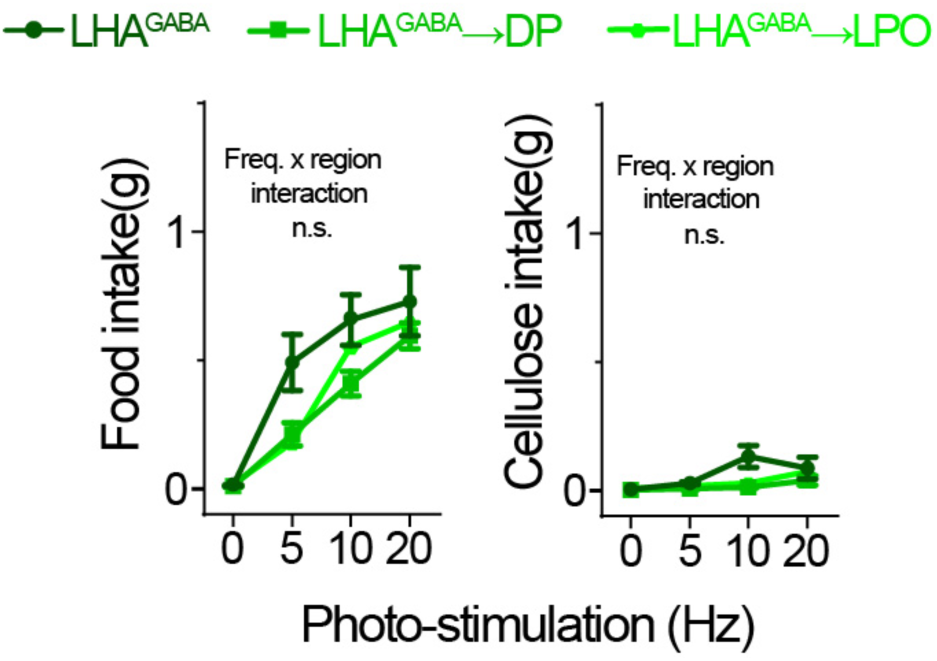
Food and cellulose intake are similar across ChR2-expressing LHA^GABA^, LHA^GABA^→DP, and LHA^GABA^→LPO mice during photo-stimulation. Two-way RM ANOVAs revealed no frequency x region interaction for food intake (*F* _4.690, 89.12_= 1.739, *P*= 0.1383) or cellulose intake (*F* _3.497, 66.44_= 2.447, *P*= 0.0625). Significant main effects of frequency were observed for food intake (*F*_2.345, 89.12_= 66.67, *P*<0.0001) and cellulose intake (*F*_1.749, 66.44_= 8.140, *P*=0. 0011). No main effect of region was observed for food intake (*F*_2, 38_= 3.125, *P*=0. 0554), whereas cellulose intake showed a marginal effect of region (*F*_2, 38_=3. 652, *P*=0. 0354). Data are mean ± SEM. Sample sizes: LHA^GABA^ (*n*=12), LHA^GABA^→DP (*n*=15), and LHA^GABA^→LPO (*n*=14). Full ANOVA results are provided in Supplemental Material.

**Figure S3.**
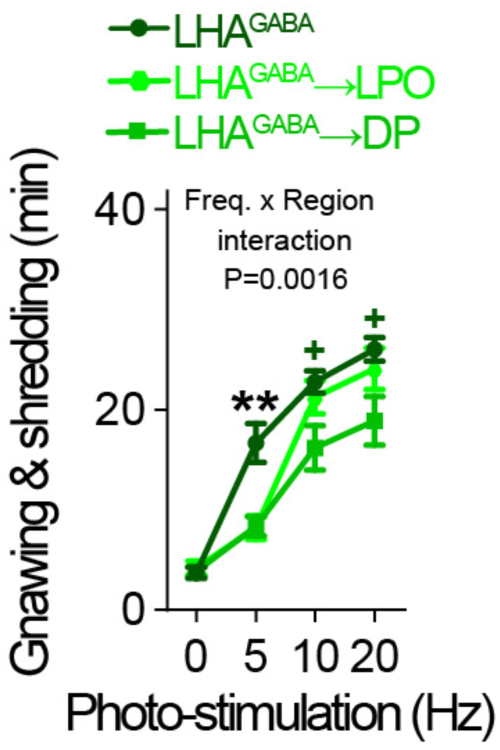
Gnawing and shredding behavior is greater following LHA^GABA^ somatic stimulation than projection-targeted stimulation. ChR2-expressing LHA^GABA^ mice showed increased gnawing and shredding compared with LHA^GABA^→DP and LHA^GABA^→LPO mice during 5-Hz photo-stimulation and compared with LHA^GABA^→DP mice during 10- and 20-Hz photo-stimulation. Two-way RM ANOVA revealed a significant frequency x region interaction (*F* _4.692, 89.14_= 4.410, *P*=0.0016), as well as main effects of frequency (*F* _2.346, 89.14_=160.2, *P*<0.0001) and region (*F*_2, 38_= 5.231, *P*=0.0098). Bonferroni post hoc tests: \*\**P*<0.01 indicate differences between LHA^GABA^ and both LHA^GABA^ → DP and LHA^GABA^→LPO groups; ^+^*P*<0.05 indicate differences between LHA^GABA^ and LHA^GABA^→DP mice. Data are mean ± SEM. Sample sizes: LHA^GABA^ (*n*=12), LHA^GABA^→DP (*n*=15), and LHA^GABA^
→LPO (*n*=14).

**Figure S4.**
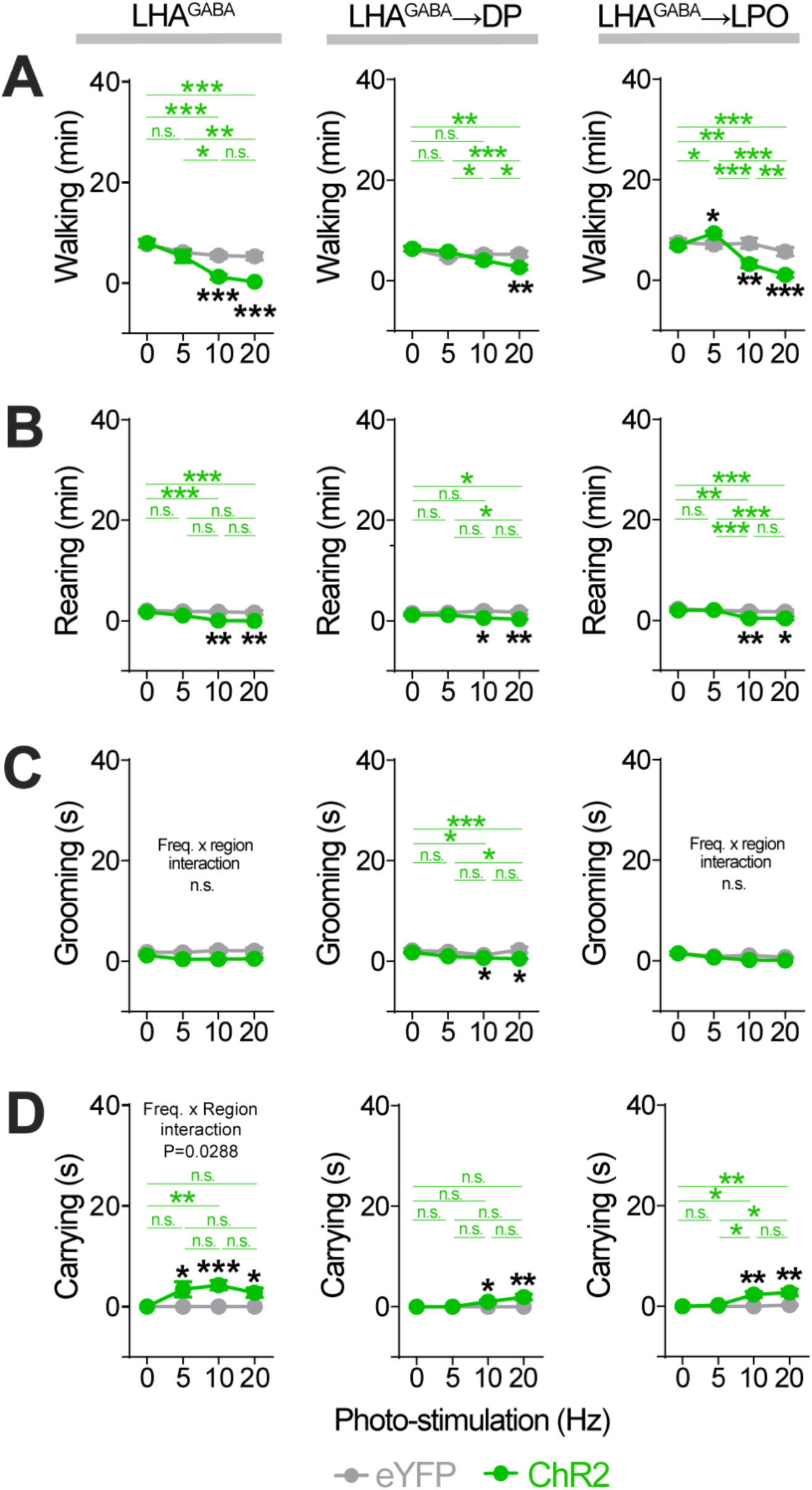
Effects of optogenetic activation of LHA^GABA^ neurons and their projections to the DP and LPO on walking, rearing, grooming, and carrying. **A** Walking time. Two-way RM ANOVA revealed significant frequency x opsin interactions for LHA^GABA^ (*F*_2.217, 46.55_= 10.15, *P*=0.0001), LHA^GABA^→DP (*F*_1.999, 43.98_=6.078, *P*=0.0047), and LHA^GABA^→LPO (*F*_2.804, 64.49_=24.08, *P*<0.0001). **B** Rearing time. Two-way RM ANOVA revealed significant frequency x opsin interactions for LHA^GABA^ (*F*_2.209, 46.39_= 5.757, *P*=0.0046), LHA^GABA^→DP (*F*_1.769, 38.91_=5.987, *P*=0.0071), and LHA^GABA^→LPO (*F*_2.257, 51.91_=5.191, *P*=0.0067). **C** Grooming time. Two-way RM ANOVA revealed a significant frequency x opsin interaction for LHA^GABA^→DP (*F*_2.665, 58.63_=3.194, *P*=0.0350), but not LHA^GABA^ (*F*_2.429, 51.01_= 2.090, *P*= 0.1247) or LHA^GABA^→LPO (*F*_1.612, 37.07_=2.179, *P*=0.1363). **D** Carrying time. Two-way RM ANOVA revealed significant frequency x opsin interactions for LHA^GABA^→DP (*F*_1.537, 33.80_=3.960, *P*=0.0383) and LHA^GABA^→LPO (*F*_2.010, 46.22_=7.067, *P*=0.0021), and trends for LHA^GABA^ (*F*_2.191, 46.01_=3.031, *P*=0.0536). Bonferroni post hoc: \*\*\**P*<0.001, \*\**P*<0.01, \**P*<0.05. Black asterisks indicate ChR2 vs. eYFP comparisons; green asterisks indicate frequency-dependent effects within ChR2 mice; n.s., non-significant. Data are mean ± SEM. Sample sizes: LHA^GABA^ (eYFP, *n*=11; ChR2 *n*=12), LHA^GABA^→DP (eYFP, *n*=9; ChR2, *n*=15), and LHA^GABA^ →LPO (eYFP, *n*=11; ChR2, *n*=14). Full ANOVA results are provided in Supplemental Material.

**Figure S5.**
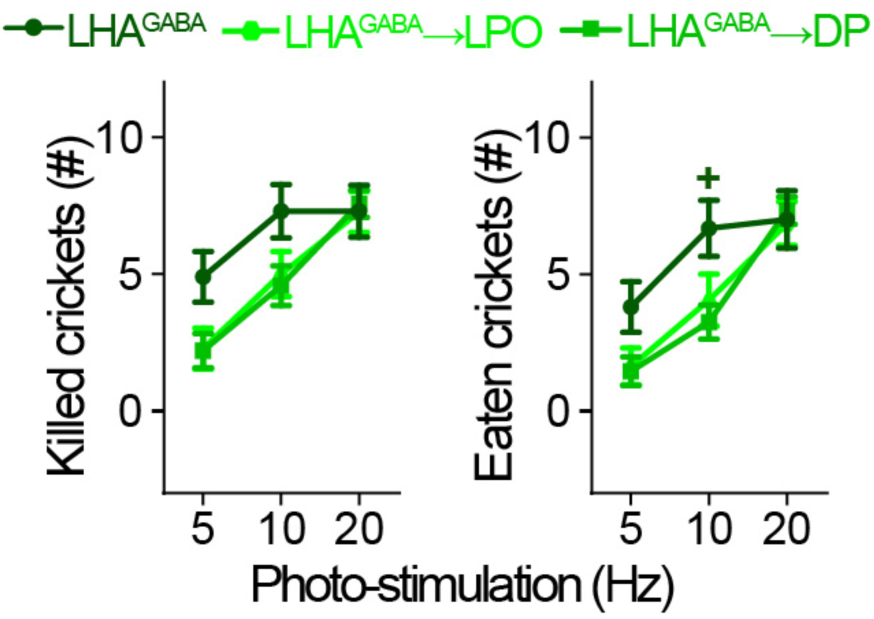
Hunting behavior is reduced in ChR2-expressing LHA^GABA^ → DP mice relative to LHA^GABA^ mice during 10-Hz photo-stimulation. *Left.* Number of crickets killed. Two-way RM ANOVA revealed a significant frequency x region interaction (*F*_3.6, 65_ =4.2, *P*=0.006) and a significant main effect of frequency (*F*_1.8, 65_=75, *P*<0.001), but no main effect of region (*F*_2, 36_= 1.7, *P*=0.200). *Right.* Number of crickets eaten. Two-way RM ANOVA revealed a significant frequency x region interaction (*F*_3.9, 70_ =5.4, *P*<0.001) and a significant main effect of frequency (*F*_2.0,70_=102, *P*<0.001), but no main effect of region (*F*_2, 36_= 1.7, *P*=0.191). Bonferroni post hoc tests indicated a significant difference between LHA^GABA^ and LHA^GABA^→DP mice at 10-Hz (^+^*P*<0.05). Data are mean ± SEM. Sample sizes: LHA^GABA^ (*n*=9) LHA^GABA^→DP (*n*=15), LHA^GABA^→LPO (*n*=14).

**Figure S6.**
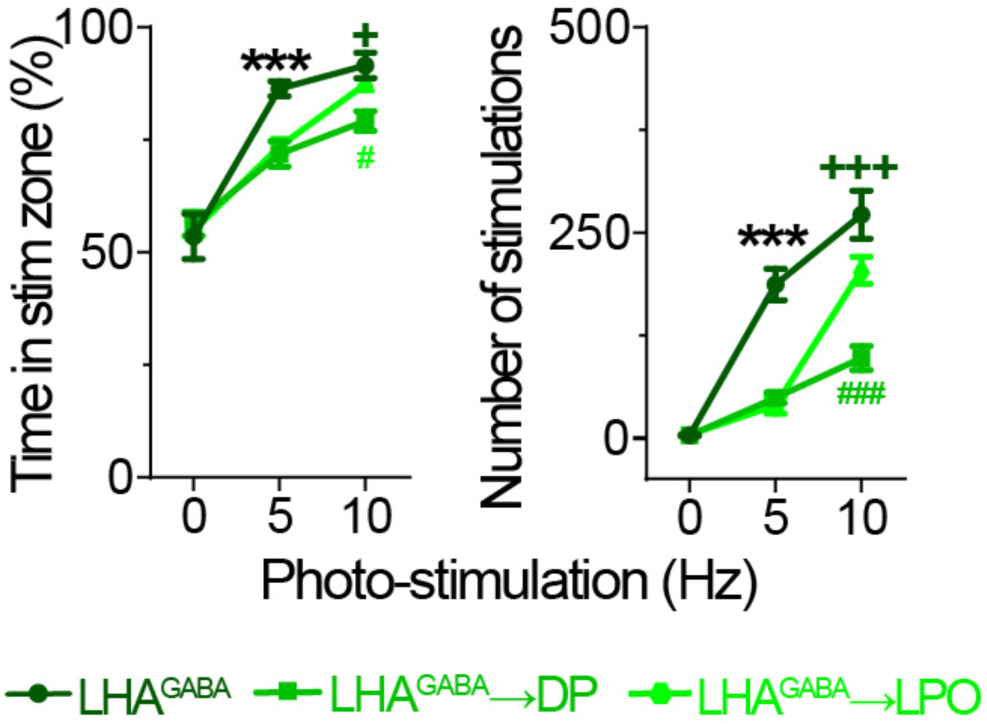
Real-time place preference and self-stimulation differ following LHA^GABA^ somatic versus projection-targeted stimulation. *Left.* Time spent in the stimulation zone (%). Two-way RM ANOVA revealed a significant frequency x region interaction (*F* _3.523, 63.41_= 5.009, *P*=0.0022), with significant main effect of frequency (*F* _1.761, 63.41_=147.7, *P*<0.0001), with significant main effect of region (*F* _2, 36_=4.760, *P*=0.0147). *Right.* Number of stimulations. Two-way RM ANOVA revealed a significant frequency x region interaction (*F* _2.846, 49.09_=18.53, *P*<0.0001), with significant main effects of frequency (*F*_1.423, 49.09_=160.3, *P*<0.0001) and region (*F*_2, 35_= 44.77, *P*<0.0001). Bonferroni post hoc tests: \*\**P*<0.01 indicates differences between LHA^GABA^ and both LHA^GABA^→DP and LHA^GABA^→LPO groups; ^+^*P*<0.05 indicates differences between LHA^GABA^ and LHA^GABA^ → DP groups; ^###^*P*<0.001, ^#^*P*<0.05 indicate differences between LHA^GABA^→DP and LHA^GABA^→LPO groups. Data are mean ± SEM. Sample sizes—RTPP: LHA^GABA^ (*n*=10), LHA^GABA^→DP (*n*=15), LHA^GABA^→LPO (*n*=14). Sample sizes—SS: LHA^GABA^ (*n*=10) LHA^GABA^→DP (*n*=15), LHA^GABA^→LPO (*n*=13).

**Figure S7.**
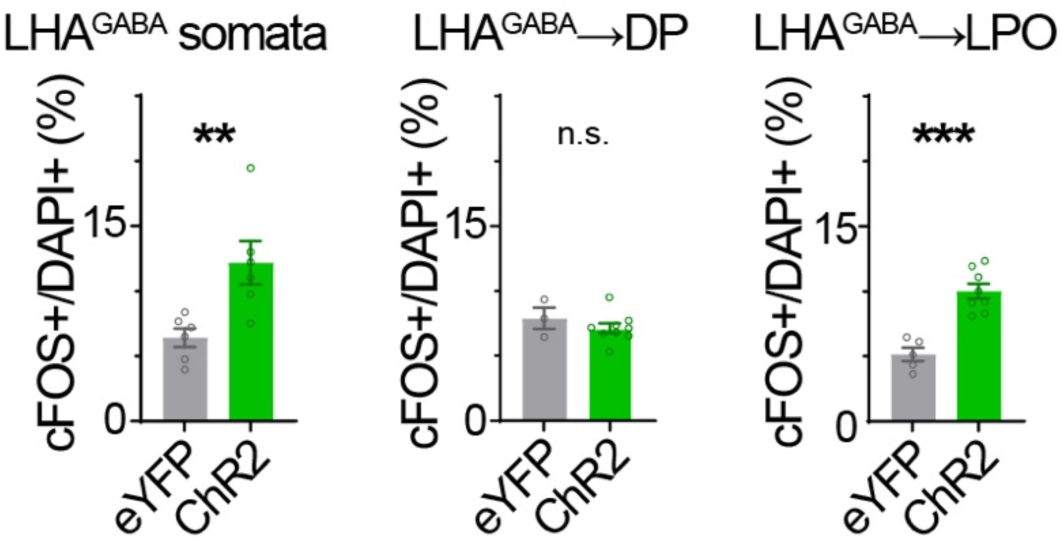
Increased LHA soma activation following stimulation of LHA^GABA^ somata and LHA^GABA^→LPO terminals in the LPO, but not LHA^GABA^→DP terminals. Percentage of Fos+/DAPI+ double-labeled somata in the LHA of ChR2-expressing LHA^GABA^ (top), LHA^GABA^→DP (middle), and LHA^GABA^→LPO mice (bottom), and their respective eYFP controls. Unpaired *t* tests: LHA^GABA^ (*t*_10_=3.191, \*\**P*=0.0096), LHA^GABA^→DP (*t*_10_=0.9552, *P*=0.3620), and LHA^GABA^→LPO mice (*t*_11_=5.803, \*\*\**P*<0.0001). Sample sizes: LHA^GABA^ (eYFP, *n*=6; ChR2, *n*=6), LHA^GABA^ → DP (eYFP, *n*=3; ChR2, *n*=9), and LHA^GABA^→LPO (eYFP, *n*=5; ChR2, *n*=8). n.s., non-significant.

## Notes

### Competing Interest Statement

The authors have declared no competing interest.

